# SAD-dependent thylakoid lipid desaturation and FDX5-associated electron transfer during copper deficiency in *Chlamydomonas reinhardtii*

**DOI:** 10.64898/2026.05.22.727179

**Authors:** Jaruswan Warakanont, Stefan Schmollinger, Samuel O. Purvine, Carrie D. Nicora, Christoph Benning, Daniela Strenkert

**Author notes:** Corresponding author: Daniela Strenkert.

## Abstract

Photosynthetic membranes undergo structural remodeling in response to environmental stress by altering fatty acid composition and desaturation levels. These changes, mediated by fatty acid desaturases (FADs), are essential for maintaining photosynthetic performance and adaptation. In this study, we demonstrate that copper-deficient *Chlamydomonas reinhardtii* cells upregulate the expression of the gene encoding stearoyl-ACP desaturase (SAD/FAB2). We propose that this four-fold induction reflects an increased physiological demand for its primary product, oleic acid (18:1*^Δ9^*), and its subsequent downstream derivatives. The *sad* mutants exhibit a significant reduction in 18:1*^Δ9^* content compared to wild-type cells, which correlates with diminished growth rates. Although SAD abundance increases under Cu deficiency, loss of SAD strongly alters C18 fatty acid composition across Cu conditions, while the growth defect is most apparent under Cu-replete conditions. This suggests that SAD activity may be a limiting factor in copper-depleted environments, leading to slower growth and reduced 18:1*^Δ9^* levels in the uncharged galactolipids monogalactosyldiacylglycerol (MGDG) and digalactosyldiacylglycerol (DGDG), both of which are critical for photosynthetic function. The desaturation reaction catalyzed by SAD requires molecular oxygen and electrons supplied by ferredoxin (Fd). Using reciprocal IP-MS, we identified FDX5 as a Cu-deficiency specific SAD interacting protein. However, *fdx5* mutants retained wild-type fatty acid profiles, indicating that FDX5 is not strictly required for SAD-dependent lipid desaturation and that another ferredoxin, likely FDX1, can compensate.

## INTRODUCTION

Fatty acids are fundamental components of bioenergetic membranes in all living organisms, with the level of fatty acid desaturation dictating membrane fluidity. Thylakoids, which are internal membranes in chloroplasts and cyanobacteria, consist of four major lipids. The most prominent lipid class within thylakoids is monogalactosyldiacylglycerol (MGDG, 52%), followed by digalactosyldiacylglycerol (DGDG, ∼27%) and to a lesser extent sulphoquinovosyldiacylglycerol (SQDG, 15%) and phosphatidylglycerol (PG, 3%) (Murphy, 1986; Webb and Green, 1991; Dörmann and Benning, 2002; Kirchhoff et al., 2002). Plant and algal thylakoid membranes have a relatively high degree of fatty acid unsaturation. Fatty acid desaturase (FAD) enzymes are responsible for modulating the level of membrane desaturation, by introducing double bonds into fatty acid hydrocarbon chains and thereby affecting membrane rigidity and affinity for individual membrane proteins. Being able to influence physiological properties of fatty acids positions FADs as key players in adjusting thylakoid membranes during adaptation to environmental stress (Wallis et al., 2002).

Three unrelated groups of FADs can be distinguished: membrane-bound FADs, soluble acyl-CoA desaturases and soluble acyl–acyl carrier protein (acyl-ACP) desaturases (Ohlrogge and Browse, 1995; Los and Murata, 1998; Sperling et al., 2003). Acyl–ACP desaturases, also known as stearoyl-ACP desaturase or Δ-9 stearate desaturase (SAD/FAB2), are soluble proteins that form homodimers and are exclusively found in the stroma of plant and algal plastids (Ohlrogge and Browse, 1995; Lindqvist et al., 1996). The fatty acid desaturation reaction catalyzed by acyl–ACP desaturases is dependent on oxygen availability and the delivery of two electrons from ferredoxin. Ferredoxins are small Fe-S cluster containing protein electron carriers, that pick up electrons resulting from oxygenic photosynthesis at photosystem I and distribute them to various client proteins in the chloroplast (Hanke and Mulo, 2013). Direct, *in vivo* evidence for ferredoxin docking to an acyl–acyl carrier protein (acyl-ACP) desaturase however has not been established. Plants and algae require constitutive production of the plastid stearoyl-ACP-Δ9-desaturase for insertion of the first *cis*-Δ9-double bond present in oleic acid (18:1*^Δ9^*, number of all carbons:number of double bonds, with Δ indicating the position of the double bond counted from the carboxyl end) in saturated fatty acids like stearic acid (18:0). This is likely an essential pathway leading to the biosynthesis of not only oleic acid but also its polyunsaturated derivatives found in all compartments of the cell (Ohlrogge and Browse, 1995). Since SADs are the only desaturases capable of utilizing the 18:0 substrate in phototrophs and the entry enzyme for the fatty acid desaturation pathway, studying how SAD activity is modulated is particularly interesting due to the enzyme’s crucial role in balancing the ratio between saturated and unsaturated fatty acids (Lindqvist et al., 1996) and thus its implications when it comes to a cell’s stress adaptation strategies by changing membrane fluidity.

*Chlamydomonas reinhardtii* is a eukaryotic alga in the green lineage (Viridiplantae), frequently used as a model system for eukaryotic photosynthesis, lipid metabolism and stress acclimation (Salomé and Merchant, 2019). Overexpression of the *FAB2* gene, which encodes the Acyl–ACP desaturase SAD, has been shown to increase oleic acid content in *Chlamydomonas reinhardtii* (Hwangbo et al., 2014). Studies of the starchless Chlamydomonas mutant BAFJ5, which exhibits increased TAG production under nitrogen starvation (Zabawinski et al., 2001), demonstrate that TAGs can be further enriched with stearic acid when grown at higher temperatures and when SAD activity was impaired (de Jaeger et al., 2017).

Curiously, *FAB2* expression is also induced when Chlamydomonas cells are grown in medium that lacks Cu supplementation (Castruita et al., 2011). This observation is concomitant with other, subtle changes to the photosynthetic apparatus under this condition. Cu is a crucial component of electron transport chains, both, in the chloroplast, from the cytochrome *b*_6_*f* complex to photosystem I (PSI), to plastocyanin (PC), and in mitochondria, at cytochrome *c* oxidase (COX2A/B). Together with the multicopper ferroxidase FOX1, which is highly abundant in Fe-limited conditions, the 3 proteins together are holding >75% of the cellular protein-bound Cu and reflect >95% of cuproprotein-encoding mRNAs (Castruita et al., 2011; Merchant et al., 2020).

Chlamydomonas grows well and performs photosynthesis even when Cu is scarce, mostly because it can substitute the soluble electron carrier plastocyanin with the heme (Fe) containing cytochrome *c*_6_ (Cyt. *c*_6_). This replacement of plastocyanin by cytochrome *c*_6_ as alternative electron carrier between the cytochrome *b*_6_*f* complex and photosystem I (PSI) in the thylakoid lumen is a well-studied example of Cu sparing, the replacement of an abundant Cu protein with a non Cu-containing alternative to preserve the available Cu for cuproenzymes that do not have a Cu-free substitute. Cu-deficient Chlamydomonas cells maintain moderate cytochrome *c* oxidase activity in the mitochondria, for which there is no Cu free substitute, by sequestering the freed Cu from plastocyanin towards this enzyme (Kropat et al., 2015).

In this work, we show that SAD protein abundance is increased by 4-fold in Cu deficient grown Chlamydomonas cells and that, as expected, 18:1 desaturation is dependent on SAD. Chlamydomonas *sad* mutants show a growth defect and accumulate their substrate in membrane lipids, stearic acid, while subsequent products are depleted. In addition, Chlamydomonas utilizes a specialized, ferredoxin isoform (FDX5), for which the gene is highly induced in response to Cu-deficiency, interacting directly with SAD in copper deficiency for electron supply. Its function in electron delivery is not essential, and can be replaced by another ferredoxin, perhaps PetF/FDX1.

## RESULTS

### SAD is four times more abundant in Cu deficient Chlamydomonas

A large-scale transcriptomics study analyzing changes with regard to copper (Cu) supply suggested that *FAB2* (encoding SAD) is induced when cells experience poor Cu nutrition (Castruita et al., 2011). This was surprising given the likely essentiality of the protein in all conditions but suggested that Cu deficient cells may have an increased demand for fatty acid desaturation. To test the impact of the observed change in *FAB2* mRNA abundance on the accumulation of the SAD protein we analyzed the abundance of the corresponding polypeptide as a function of Cu nutrition. For this purpose, we used a commercially available antiserum targeting SAD, and in addition we also generated Chlamydomonas strains that produce SAD-HA tagged fusion proteins directly from their native locus, using CRISPR/Cpf1 mediated gene knock-in technology (Pham et al., 2022; Strenkert et al., 2024). In brief, algal cells were transformed by electroporation in the presence of the Cpf1/gRNA ribonucleoprotein complex targeting the *FAB2* gene and a single stranded DNA repair template (ssODN), containing homology arms to *FAB2* and a sequence coding for a glycine linker and a codon optimized DNA sequence encoding the human influenza hemagglutinin (HA) protein (Figure S1 and Table 1). We identified two strains that incorporated the *in-frame* sequence encoding linker and the HA tag at the right position, designated SAD-HA #107 and SAD-HA #198 (Figure 1a). As expected, HA-tagged SAD proteins (including the linker sequence) exhibited a slight electrophoretic shift, migrating more slowly during SDS-PAGE than their unmodified counterparts. This shift was confirmed using an anti-SAD antiserum (Figure 1a). Furthermore, the anti-HA antibody detected a single band at the predicted

**Figure 1.**
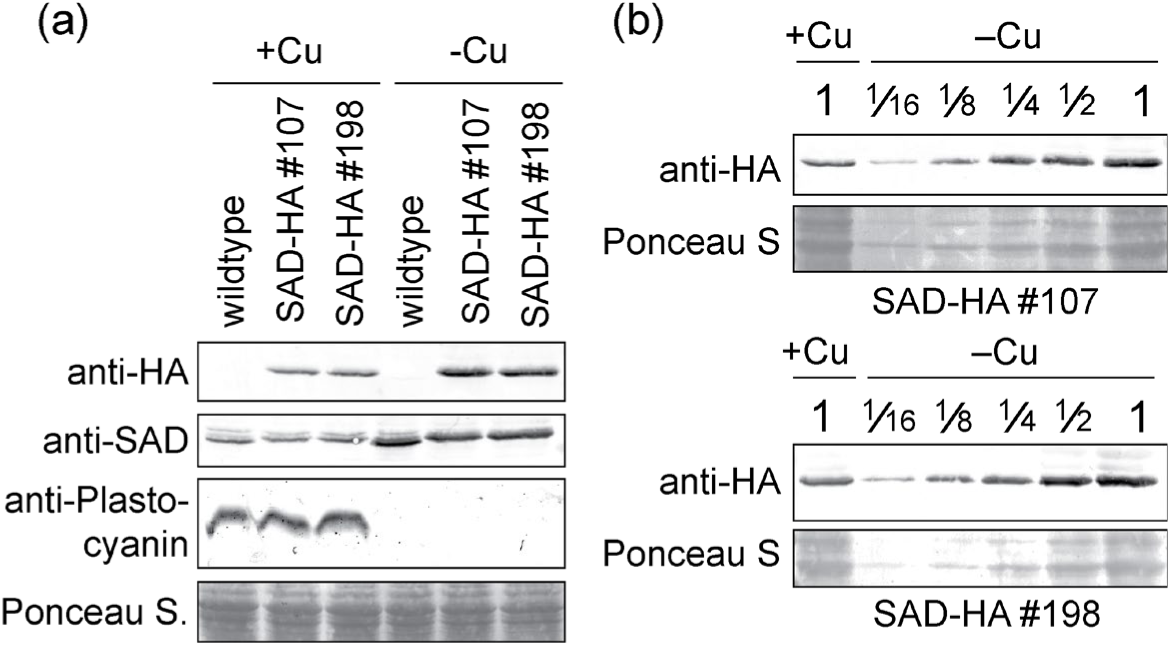
SAD abundance depends on Cu nutritional status. (a,b) *Chlamydomonas reinhardtii* wildtype, SAD-HA #107 or SAD-HA #198 strains were grown in either Cu-replete TAP medium (+Cu) or in Cu-deficient (−Cu) growth medium as indicated. Total protein was separated by 15% SDS PAGE and immunodetected using antisera against SAD, plastocyanin or HA. Ponceau S stain was used as loading control. molecular weight specifically in lysates from cells producing the SAD-HA fusion protein, whereas no signal was observed in wild-type samples (Figure 1a). Chlamydomonas wild-type and SAD-HA cultures grown in TAP medium without Cu supplementation exhibited the expected degradation of plastocyanin (Merchant et al., 2020) (Figure 1a), confirming the successful induction of Cu deficiency. Immunoblot analysis revealed that SAD protein accumulation increased 4-fold under Cu deficiency (Figure 1ab).

**Table 1.**
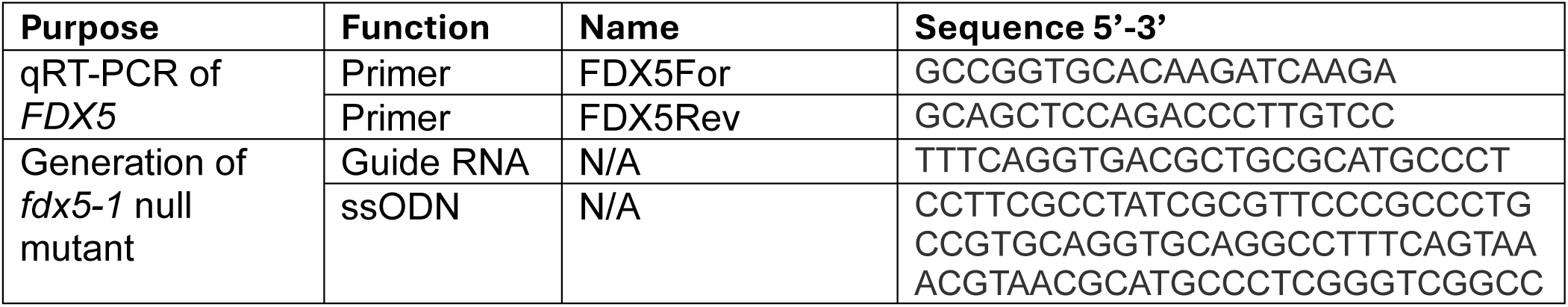
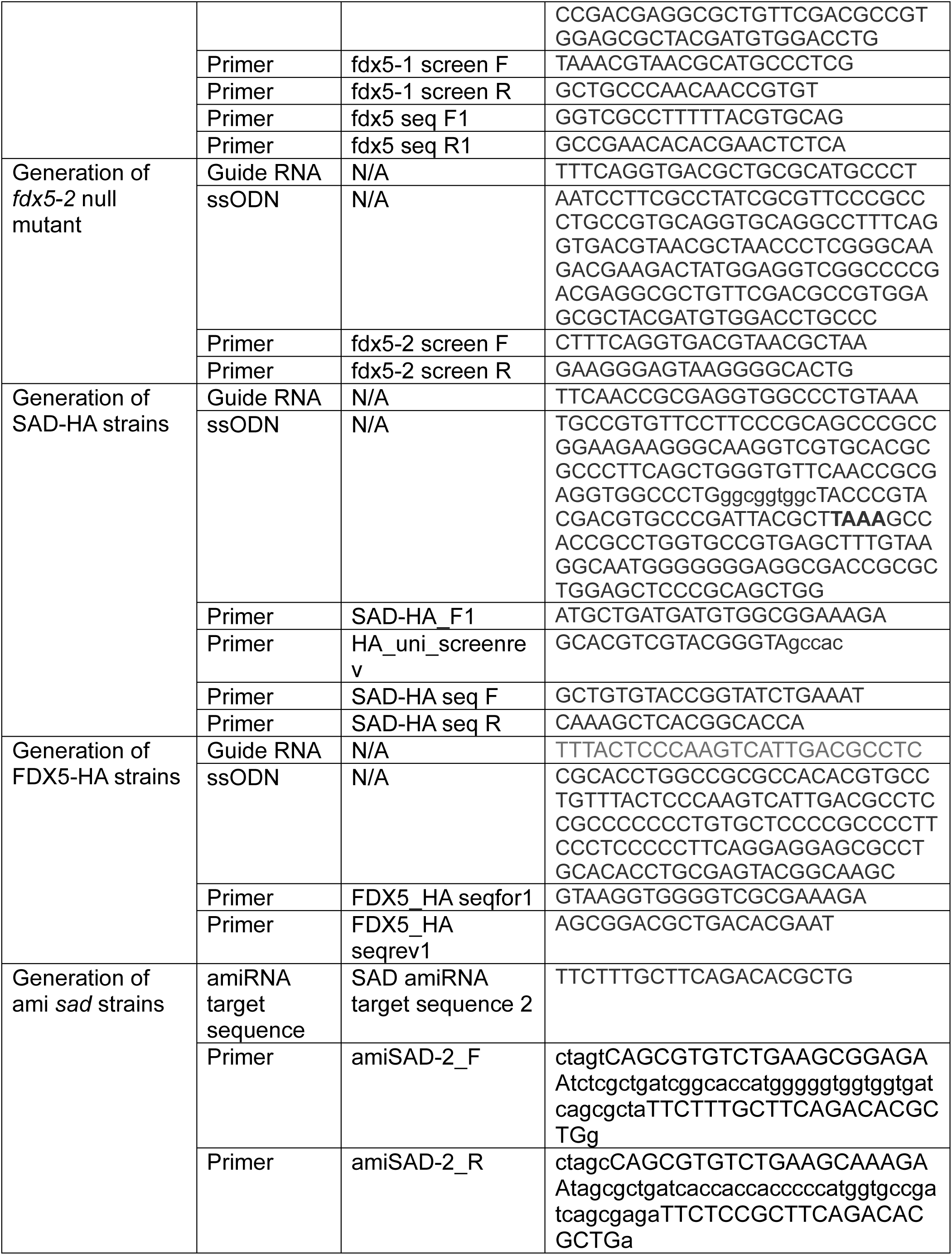
Sequences of primers and oligonucleotides for CRISPR.

These results demonstrate that SAD protein levels increase during Cu deficiency, consistent with previously reported mRNA abundance changes. However, the physiological rationale for this upregulation remains to be elucidated. We hypothesize that Cu limitation triggers an increased demand for oleic acid and its unsaturated derivatives. Specifically, a higher degree of fatty acid desaturation within the thylakoid membrane may be required to maintain membrane functionality during Cu-deficiency dependent remodeling of the bioenergetic membrane, concomitant with the substitution of the soluble electron carrier plastocyanin with cytochrome *c*_6_.

### Mutants in SAD show reduced growth and impaired oleic acid synthesis

To investigate the Cu-dependent function of SAD (encoded by *FAB2*) in Chlamydomonas, we designed an artificial microRNA (amiRNA) targeting the 3′ region of the *FAB2* gene (Figure S1) (Molnár et al., 2009; Schmollinger et al., 2010). We initially screened candidate *sad* knockdown lines for a reduced 18:1*^Δ9^*to 18:0 fatty acid ratio relative to wild-type (wt) strains transformed with an empty vector control. Consistent with our lipid screening results, we identified three lines with clearly reduced SAD protein levels (Figure 2a), which we designated ami-*sad1*, ami-*sad2*, and ami-*sad3*. Compared to the empty-vector control strains, SAD protein abundance was reduced to less than 10% in all three ami-sad lines, regardless of whether they were grown in Cu-supplemented (+Cu) or Cu-deficient (-Cu) medium. Among these, ami-*sad1* represented the weakest knockdown, while ami-*sad3* exhibited the most significant reduction in SAD protein levels (Figure 2a). If SAD-mediated remodeling of chloroplast membrane lipids is essential for accommodating structural changes to the photosynthetic machinery during Cu deficiency, *sad* knockdown strains should exhibit impaired photosynthetic performance and, consequently, a discernible growth phenotype when grown in Cu-deficient medium. To test this hypothesis, we cultivated the ami-*sad* lines alongside the corresponding wt strains in both Cu-replete and Cu-deficient growth medium.

**Figure 2.**
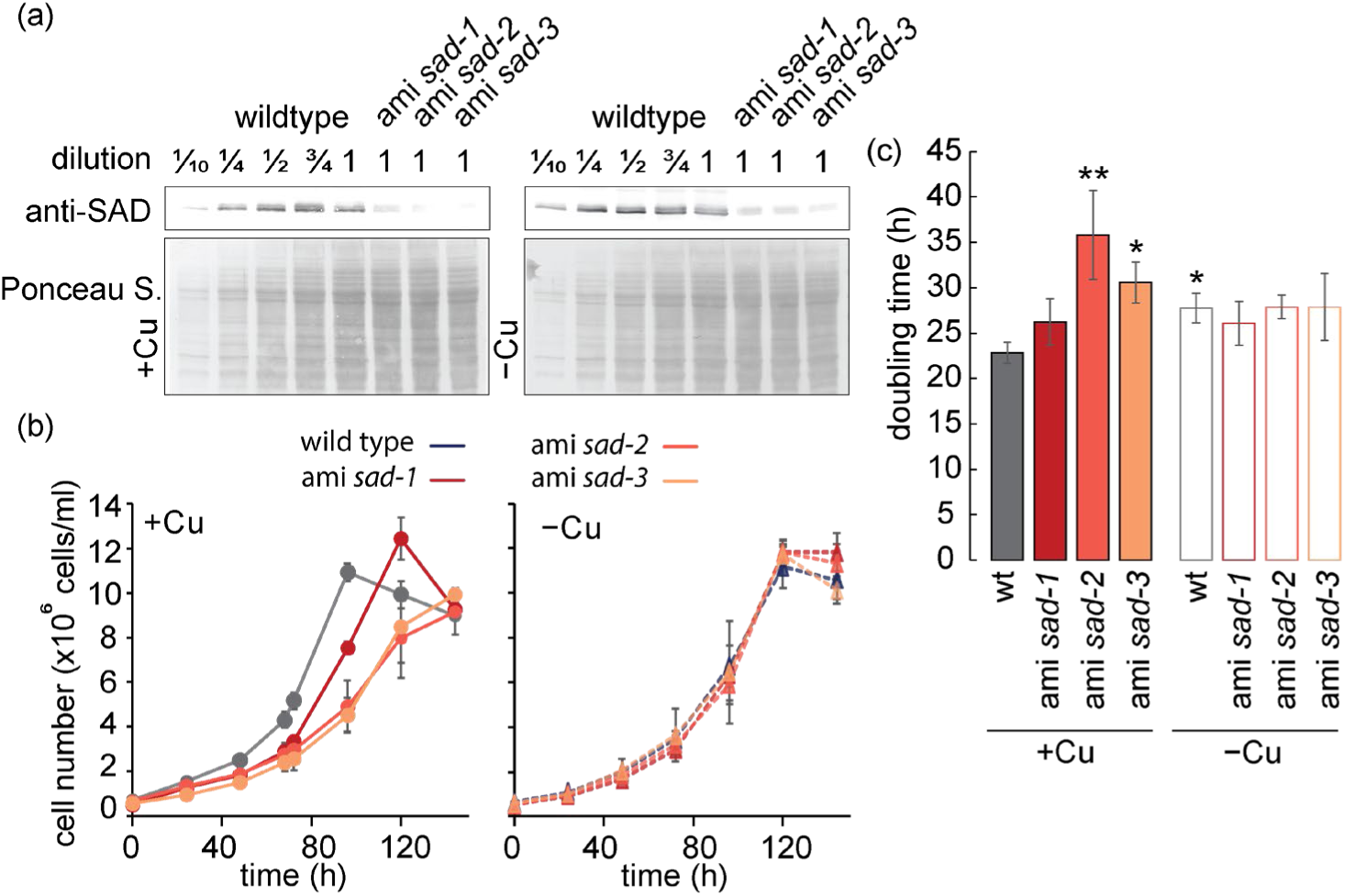
Mutants with reduced SAD protein show reduced growth that is rescued by Cu deficiency. Wildtype strains (grey), ami *sad-1* (red), ami *sad-2* (dark orange) and ami *sad-3* (orange) were grown in either Cu-replete TAP medium (+Cu) or in Cu-deficient (−Cu) growth medium as indicated. (a) Total protein was separated by 15% SDS PAGE and immunodetected using antisera against SAD, plastocyanin or HA. Ponceau S stain was used as loading control. (b) Cell counts were obtained at 24-h intervals. Shown are averages and standard deviation from three independently grown cultures. (c) Doubling times were calculated from data obtained in (b) during exponential growth. Data are means ± SD from three independently grown cultures. Asterisks indicate significant differences compared to wt (+Cu) (two-sided *t*-test, P < 0.05).

Under Cu-replete conditions, the *ami-sad* lines exhibited reduced growth rates compared to wt cells. The severity of this growth defect correlated with the degree of SAD knockdown, with the ami-*sad2* and ami-*sad3* mutants showing the most significant growth impairment (Figure 2b). Wt cells grown under Cu-deficiency exhibited slower growth compared to copper-replete cultures, with doubling times increasing from 23 to 28 hours (Figure 2c). Interestingly, no significant difference in growth was observed between ami-*sad* mutants and wild-type strains under Cu deficiency.

Previous studies have demonstrated that Cu deficiency alters the composition of chloroplast membrane lipids by increasing the degree of fatty acid desaturation (Castruita et al., 2011). This shift is accompanied by the transcriptional up-regulation of genes coding for multiple fatty acid desaturases (FADs) under -Cu conditions, including S*AD*, the ω-6 fatty acid desaturase (*FAD6*) and the MGDG-specific palmitate Δ-7 desaturase (*FAD5a*). To determine which fatty acid alterations are lipid-class specific and dependent on SAD activity, we purified individual lipid fractions for analysis. Comparisons were made between ami-*sad* and wt strains grown in media either supplemented with or depleted of Cu. As anticipated, oleic acid (18:1*^Δ9^*) biosynthesis was impaired in the *ami-sad* lines regardless of copper availability. This reduction in 18:1*^Δ9^* occurred concomitantly with an accumulation of the 18:0 precursor (Figure 3a). The levels of several polyunsaturated fatty acids (18:2*^Δ9,12^* and 18:3*^Δ5,9,12^*) were also reduced in the *ami-sad* mutants. Similar to the observations for 18:1*^Δ9^*, these reductions occurred independently of the copper concentration in the growth medium (Figure 3a). C_16_ fatty acids were largely unchanged in ami-*sad* mutants. These significant alterations in C18 fatty acids were observed in both monogalactosyldiacylglycerol (MGDG) and digalactosyldiacylglycerol (DGDG). These uncharged galactolipids are essential components of the thylakoid membrane and play crucial roles in maintaining photosynthetic function. MGDG was the primary lipid species found to increase in response to Cu deficiency. This accumulation was maintained in the *sad* mutants (Figure S2), suggesting a heightened demand for this lipid class under these conditions. In galactolipid fractions (MGDG and DGDG), the ami-*sad* strains also exhibited increased saturation, specifically, elevated 18:0 alongside reduced 18:1*^Δ9^* and 18:2*^Δ9,12^* levels compared to the wt (Figure 3bc). These results demonstrate that the loss of SAD activity alters the fatty acid profile of the thylakoid membrane. 18:2*^Δ9,12^*is one of the main fatty acids increasingly accumulating in Cu deficient cells, both globally and in galactolipid fractions (Figure 3abc). Unexpectedly, the decrease in 18:1*^Δ9^* and 18:2*^Δ9,12^* fatty acids did not result in a decrease in 18:3*^Δ9,12,15^*, the major product of 18:2*^Δ9,12^* desaturation (Figure 3b). 18:3*^Δ9,12,15^* is one of the two major fatty acids in MGDG, it was still accumulating to at least wildtype levels, indicating that its desaturation from 18:2*^Δ9,12^* was not affected by SAD reduction (Figure 3b). In addition to MGDG accumulation, both wt and ami-*sad* strains sequestered triacylglycerols (TAG) under Cu-deficient conditions (Figure S2). This finding is consistent with the observed formation of lipid bodies in a Cu deficient cell (Kropat 2011).

**Figure 3.**
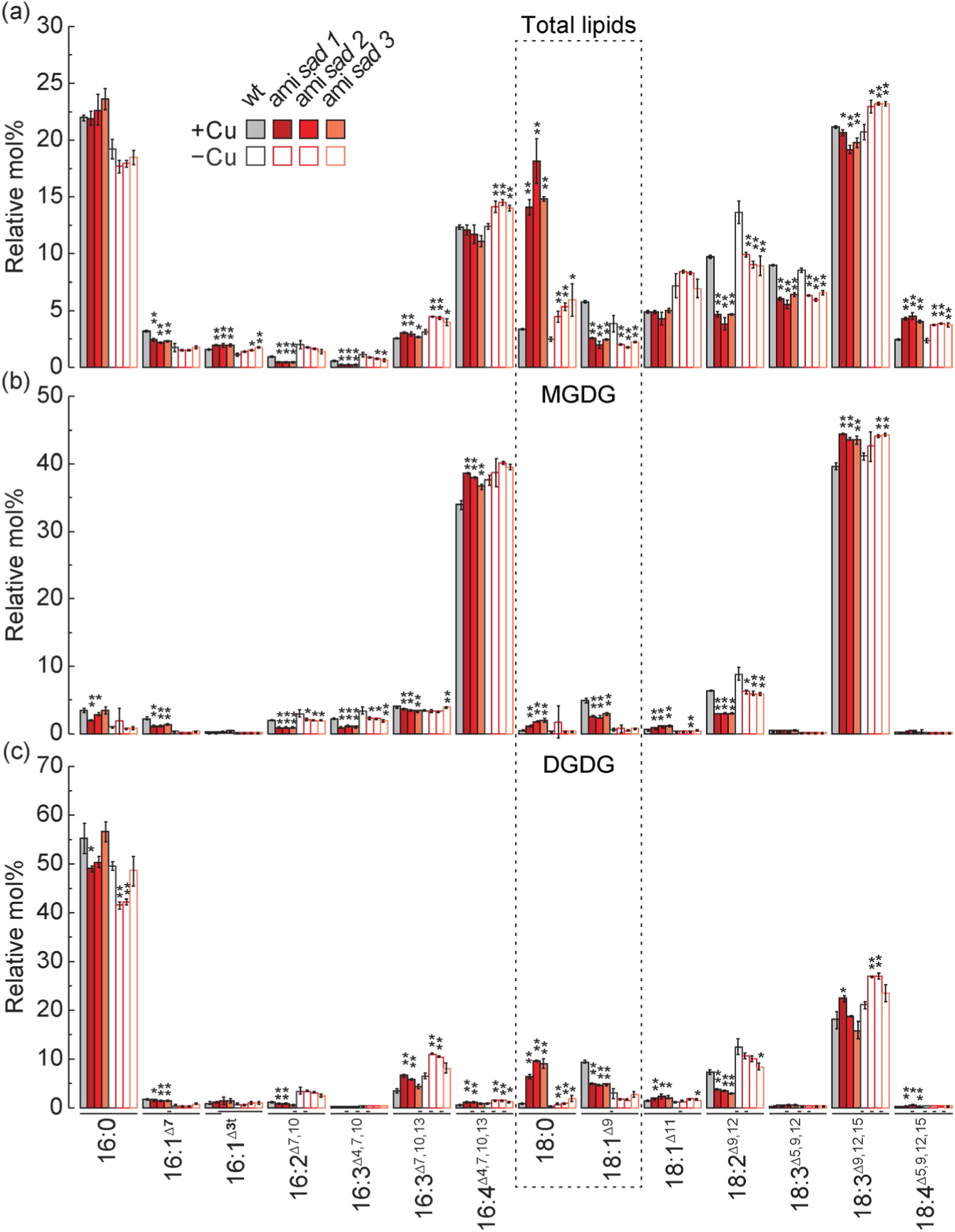
**Changes in acyl group profiles in ami *sad* mutants**. Acyl group profile of total lipids (a), monogalactosyldiacylglycerol (MGDG, b) and digalactosyldiacylglycerol (DGDG,c) of the wt and ami *sad* mutants during mid-logarithmic phase grown in different copper (+Cu and −Cu conditions. The bars show fatty acid composition in mol % and indicate the means (±SD) of three independent experiments. Standard nomenclature for fatty acids is used, and is indicated below the *x*-axis in (c): number of carbons:number of double bonds, with position of double bonds indicated counting from the carboxyl end. A single asterisk (*) and a stacked double asterisk (one * above another *) indicate statistically significant differences from the wild type with *p* ≤ 0.05 and *p* ≤ 0.01, respectively, based on two-tailed Student’s *t*-tests assuming equal variances. Statistical analysis was first performed using one-way ANOVA among wild type and mutant groups. Pairwise *t*-tests were conducted only when the ANOVA result was significant (*p* ≤ 0.05). Stearic and oleic acid are highlighted by a grey box.

Taken together, *sad* knockdown mutants show reduced growth in Cu-replete conditions and have aberrant fatty acid composition involving 18 carbon fatty acids, resulting in higher levels of saturated fatty acids, and reduced levels of (mono/poly)-unsaturated fatty acids globally (18:1*^Δ9^*, 18:2*^Δ9,12^*, 18:3*^Δ5,9,12^*) and in galactolipids specifically (18:1*^Δ9^*, 18:2*^Δ9,12^*), largely independent of Cu nutritional status.

### The Cu nutrition-dependent SAD interactome

SAD introduces a double bond into saturated fatty acids, primarily converting stearate (18:0) into monounsaturated oleate (18:1*^Δ9^*). This oxidation reaction requires both molecular oxygen and a reductant. In Chlamydomonas, the latter is likely provided by one of the 12 ferredoxin (Fd) isoforms encoded in the genome (Schmollinger et al., 2026). To identify SAD-interacting partners, specifically the Fd isoform serving as the soluble electron carrier between Photosystem I (PSI) and SAD, we performed immunoprecipitations (IP) followed by mass spectrometry (MS). These analyses utilized two independent SAD-HA producing strains (SAD-HA #107 and #198, Figure 1) cultivated in growth medium with and without copper supplementation. A wild-type strain lacking the SAD-HA fusion protein was included as a negative control. This allowed us to filter the mass spectrometry data and identify proteins that were enriched non-specifically. For IP, cells were collected, lysed by sonication and immuno-precipitated with an anti-HA antibody. Using this approach, we immunoprecipitated 117 and 112 proteins from Cu-replete SAD-HA cultures and 137 and 100 proteins from Cu-deplete SAD-HA strains, respectively. None of these proteins were detected in the wild-type controls (Figure 4, Data S1). Among these candidates, 63 proteins were enriched in the Cu-replete samples and 36 in the Cu-deficient samples across both independent SAD-HA expressing strains (Data S1). Regardless of the Cu nutritional status, SAD was the most abundant protein identified by MS in both SAD-HA strains. Its complete absence in the wild-type immunoprecipitates confirms that the bait protein was successfully and specifically enriched (Data S1). Independent of Cu supplementation, SAD was found to interact with acyl-carrier protein 2 (ACP2, Data S1), a key component of the plastid fatty acid biosynthetic machinery. Acyl-carrier proteins (ACPs) bind saturated fatty acids and serve as the essential substrates for SAD-mediated desaturation (McKeon and Stumpf, 1982; Schultz et al., 2000).

**Figure 4.**
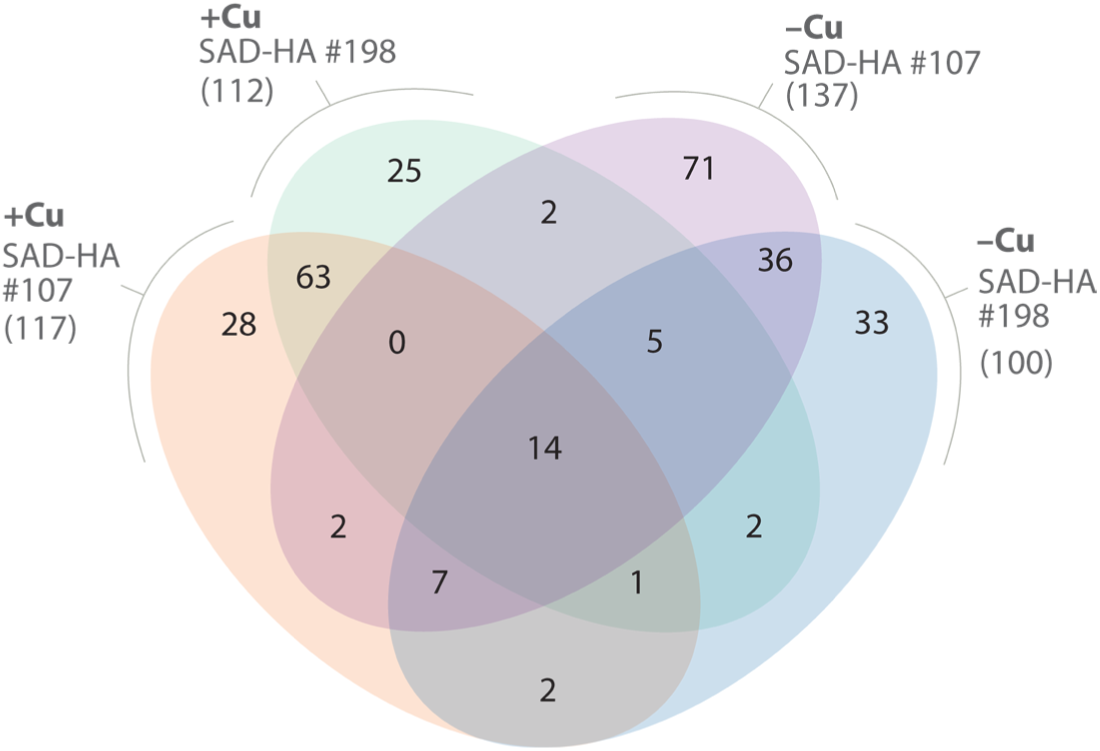
SAD interacts with ACP-carrier protein and FDX5 in copper deficiency. Venn diagram shows the overlap of proteins co-immunoprecipitated with SAD-HA under different copper (+Cu or −Cu) conditions from two different SAD-HA expressing strains (#107 and #198). Numbers in parentheses represent total numbers of protein identified in each condition. Each region of the diagrams shows the number of protein unique to or shared between specific combinations of these conditions.

While this interaction was anticipated, it has not been previously demonstrated *in vivo* in Chlamydomonas. These results validate our experimental system and demonstrate our capacity to enrich canonical SAD-interacting proteins. Notably, we identified the ferredoxin isoform FDX5 as a potential interactor in cells grown under Cu deficiency only; it was enriched in the Cu deficient SAD-HA #198 strain but absent in both the wt and the other SAD-HA strain. FDX5 is one of 12 annotated Fds in Chlamydomonas and is FDX5 is strongly induced at the transcript level under Cu deficiency (Figure S3). Crucially, FDX5 was the only Fd detected in our SAD-HA co-immunoprecipitation (CoIP) experiments. This observation was intriguing, as it implies that a specialized ferredoxin isoform, FDX5, may have evolved to supply SAD with electrons specifically in a Cu deficient cell. The recovery of Fds by mass spectrometry is often hindered by their low molecular weight and the highly transient nature of Fd-client interactions; consequently, the identification of FDX5 in our dataset is a notable finding. The identification of FDX5 is consistent with a previous transcriptomic study reporting a 192-fold induction of *FDX5* expression under Cu-deficiency (Castruita et al., 2011). This strong upregulation suggests that FDX5 plays a role in reshaping chloroplast metabolism in response to Cu limitation, a process that includes the observed, strategic remodeling of thylakoid membrane lipids.

To confirm the interaction between FDX5 and SAD, we generated two independent FDX5-HA expressing strains, FDX5-HA #25 and FDX5-HA #221 (Figure 5a, Figure S4). These strains were engineered using the same CRISPR/Cpf1-mediated approach previously employed for the SAD-HA strains (Figure S4, Table 1). While FDX5-HA #221 contained the intended gene edits, FDX5-HA #25 contained a 9-bps deletion within an upstream intron (Figure S4).

**Figure 5.**
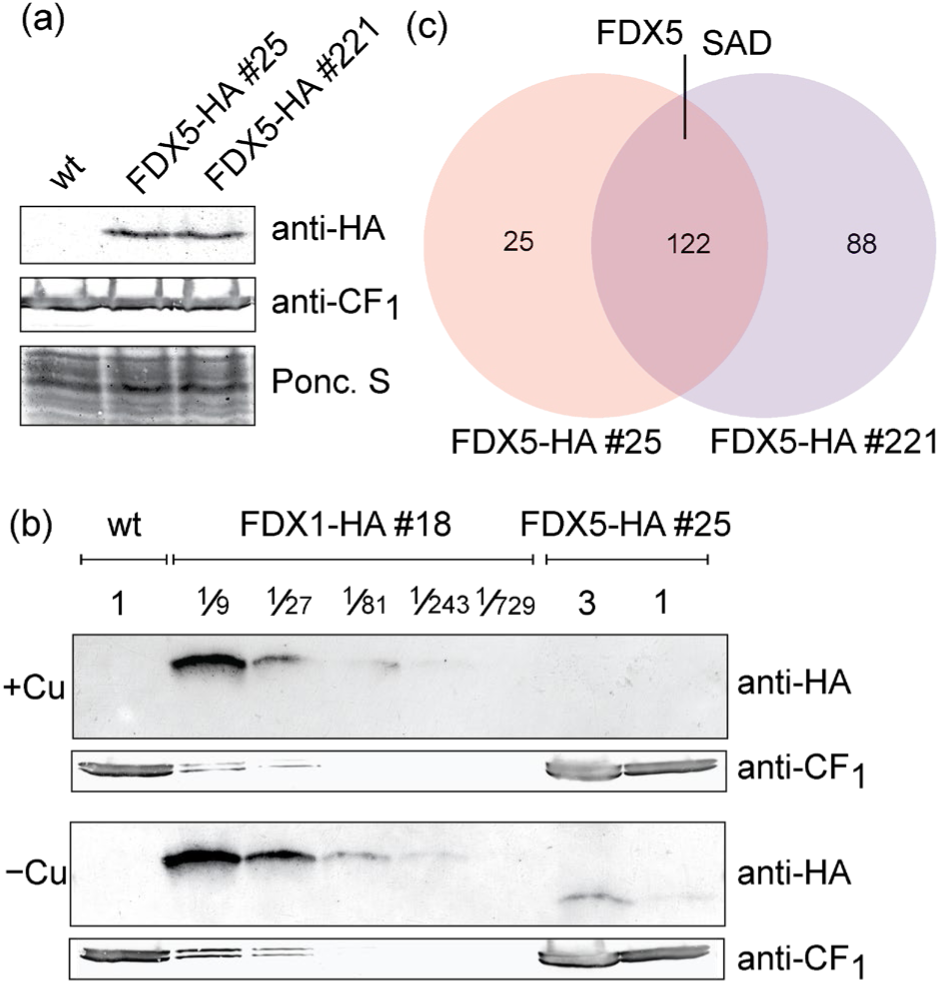
FDX5 accumulates to ∼0.4% relative to FDX1 protein and interacts with SAD in Cu deficient cells. (a) Immunoblot analysis of wt, FDX5-HA strains #25 and #221 grown under Cu deficiency. Protein samples were normalized by cell number, with extracts from 2 × 10⁶ cells loaded per lane. Equal loading of proteins was verified by Ponceau S (Ponc. S) staining (total protein) and anti-CF_1_ signal (chloroplast ATP synthase CF_1_ subunit) across lanes. (b) Immunodetection of FDX1-HA #18 and FDX5-HA #25 against anti-HA antibody. Proteins were loaded in serial dilutions; dilution 1 corresponds to protein extracted from 2 × 10⁶ cells. Anti-CF_1_ signal indicates protein loading. (c) Venn diagram shows the overlap of proteins co-immunoprecipitated with FDX5-HA under copper deficiency conditions from two different FDX5-HA expressing strains (#25 and #221). Numbers in parentheses represent total number of proteins identified in each condition. Each region of the diagram shows the number of proteins unique to or shared between the two FDX5-HA strains.

Consistent with our results for SAD-HA, the FDX5-HA protein was detected exclusively in the engineered strains. It migrated at the expected molecular weight during SDS-PAGE in both FDX5-HA expressing strains (Figure 5a), with no additional bands observed. Under Cu-deficient conditions, *FDX5* accounts for approximately 40% of the total Fd mRNA pool (Figure S3), whereas it is barely detectable (<1%) in Cu-replete strains. To estimate the abundance of FDX5, we compared its levels to those of FDX1 (PetF). FDX1 is the primary plastid Fd in Chlamydomonas and is constitutively present in high levels (Figure S3) (Schmollinger et al., 2026). FDX1 plays an essential role in photosynthesis by transferring electrons from Photosystem I to Fd-NADP^+^ reductase (FNR) for NADPH production. To determine the relative stoichiometry of FDX1 and FDX5 within the chloroplast under Cu-replete and Cu-deplete conditions, we utilized two independent FDX1-HA producing strains previously generated in our laboratory (Schmollinger et al., 2026). Given the substantial increase in *FDX5* transcript levels during Cu deficiency, we hypothesized that the FDX5 protein might accumulate to levels comparable to FDX1. However, our results did not support this. To resolve the relative protein abundances, we performed iterative comparisons using serial dilutions of whole-cell protein from FDX1-HA and FDX5-HA expressing strains. As expected, FDX5-HA was not detected in Cu-replete strains (Figure 5b). In Cu-deficient strains, FDX5 accounted for only approximately 0.4% of the FDX1 protein abundance (Figure 5b), a value roughly 100-fold lower than what would be predicted based solely on mRNA levels. Although we confirmed the copper-dependent induction of the FDX5 protein as predicted by previous transcriptomic data (Figure S3), its contribution to the total ferredoxin protein pool is far less significant than anticipated.

### FDX5 interacts with SAD but also other known Fd dependent enzymes

To validate the interaction between SAD and FDX5 and identify potential FDX5-client proteins, we performed CoIP experiments using FDX5-HA as bait. Given that SAD is a larger, soluble protein and was successfully identified in our previous reciprocal experiments, we anticipated it would be detectable by the IP-MS approach. Immunoprecipitation was performed specifically on Cu-deficient FDX5-HA and wild-type strains, as FDX5 presence is restricted primarily to these conditions (Figure 5b and Data S1). A total of 122 proteins were identified by IP-MS across both FDX5-HA strains, all of which were entirely absent in the wild-type controls (Figure 5c, Data S1). Among these candidates, we identified both the bait protein, FDX5, and its partner, SAD, thereby validating the *in vivo* interaction between these two proteins (Data S1). Acyl-carrier protein 2 (ACP2), which serves as a known substrate for SAD, was also enriched in one of the two experiments. This suggests that the pull-down successfully captured the larger SAD-ACP2-FDX5 complex. In addition to SAD, we identified the Photosystem I (PSI) reaction center subunit II (PSAD1) among the 122 proteins enriched in both FDX5-HA strains. This finding suggests that PSAD1 may serve as the primary docking site for FDX5. We also identified several other ferredoxin-dependent enzymes in the FDX5-HA pull-downs, including sulfite reductase 2 (SiR2), the iron hydrogenase HydA1, and its associated maturation proteins (HydEF). These enzymes are characteristically co-produced with FDX5 when strains are grown under Cu-limited or hypoxic conditions. This direct evidence of FDX5 interacting with Fd-dependent enzymes suggests that FDX5 has evolved unique properties to sustain reductive metabolism in Cu-deficient strains, capabilities that may be absent in other ferredoxin isoforms.

### *fdx5* mutants are asymptomatic in Cu deficiency and exhibit minor proteomic changes

The direct interaction between SAD and FDX5 under Cu deficiency suggests that FDX5 may serve as the primary electron donor for this enzyme. To investigate the role of FDX5 in fatty acid desaturation, we generated *fdx5* null alleles by introducing in-frame stop codons into the first exon using the CRISPR/Cpf1 approach described above (Figure S4 and Table 1).

Following diagnostic PCR and validation via Sanger sequencing, two strains, designated *fdx5-1* and *fdx5-2*, were identified. Both exhibited significantly reduced *FDX5* transcript levels compared to the wt, concomitant with a reduction in the corresponding polypeptide to below detection limit in mass spectrometry (Figure S5) and both mutants were selected for subsequent phenotypic characterization (Figure S5).

Growth of the *fdx5* mutants was indistinguishable from wt under both Cu-replete and Cu-limited conditions (Figure S5b). If SAD activity were exclusively dependent on FDX5 for electron delivery in Cu-deficient strains, we would expect lower levels of 18:1^Δ9^ in the *fdx5* mutants, however, this was not observed. Indeed, the fatty acid profiles of total lipids, MGDG, and DGDG showed no significant differences between the independent *fdx5* mutants and wt strains under either condition (Figure S6).

To determine whether *fdx5* mutants exhibit alternative molecular phenotypes, we conducted an untargeted survey of the proteome. We grew *fdx5* mutants and wt strains under conditions where FDX5 expression is maximally induced: Cu deficiency or dark hypoxia (Castruita et al., 2011; Hemschemeier et al., 2013). As anticipated, FDX5 was detected in wt strains under either Cu-deficient or hypoxic conditions, yet it was entirely absent in both *fdx5* mutants (Data S2, Figure 6). We captured the Cu conditional or dark hypoxic response, evident by the expected expression patterns of marker proteins cytochrome *c*_6_ (Cu deficiency marker) and Fe-hydrogenase (dark hypoxia marker), respectively (Figure 6). A total of 1544 proteins were quantified in all samples (Figure S7). We used principal component analysis (PCA) for multivariate analysis of proteome dynamics based on protein abundances. Samples were separated by condition, with wt and *fdx5* mutants clustering closely together (Figure S7).

**Figure 6.**
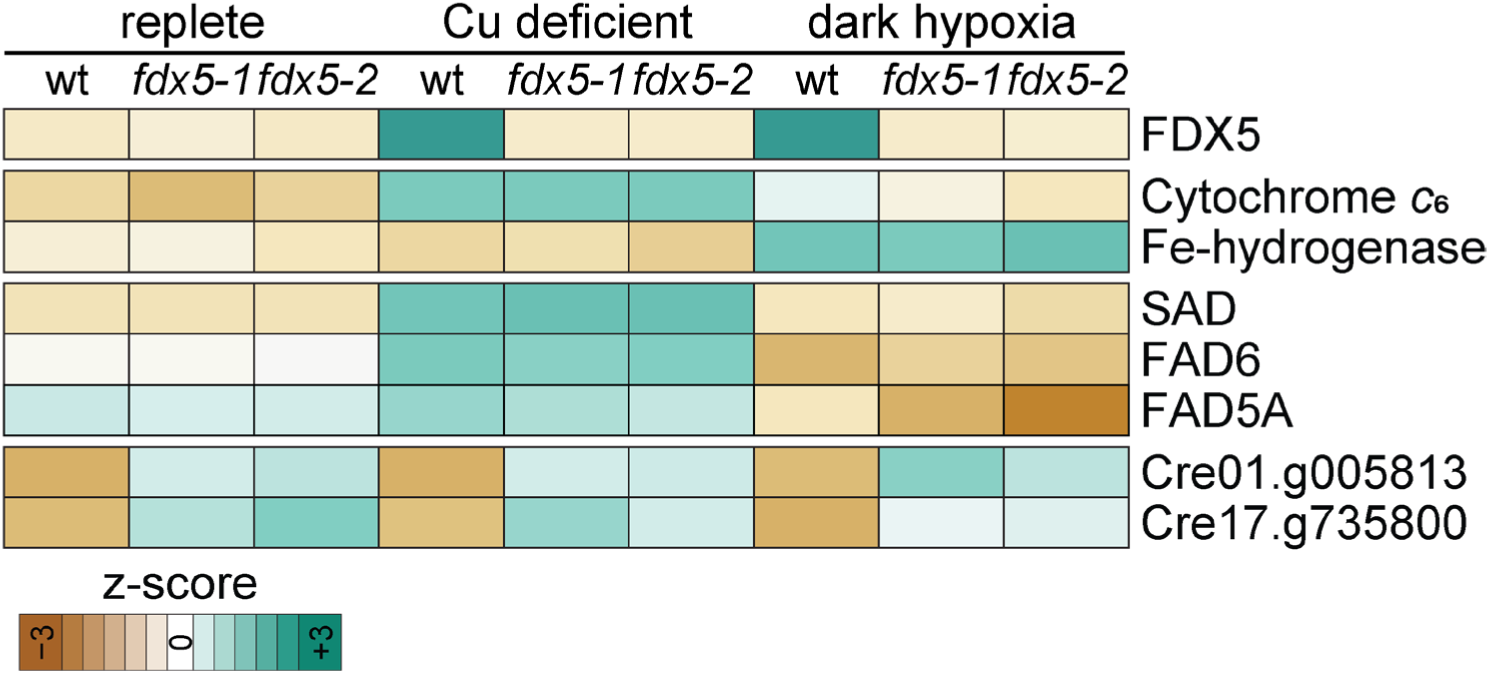
Proteomic changes in *fdx5* mutants grown under Cu deficiency or dark hypoxia. Heatmap of proteins identified in wt, *fdx5-1* and *fdx5-2* mutants. Cells were grown in medium with (replete), without Cu supplementation (Cu deficient) or for 48 h in dark hypoxia according to Hemschemeier *et al*. 2013. Protein intensities are displayed as colours ranging from brown to turquoise as shown in the key.

Consistent with the lack of a discernible growth defect, changes in the proteome were relatively minor. We identified just two proteins of unknown function that displayed significant differences in abundance between the *fdx5* mutants and wt strains under both conditions (Figure 6, Data S2). Both genes are expressed at low levels (2–4 FPKM) and remain unaffected by copper deficiency or dark hypoxia (Castruita et al., 2011; Hemschemeier et al., 2013).

These data suggest that the loss of FDX5 function may be functionally complemented by another Fd isoform, with the highly abundant FDX1 being the most likely candidate.

## DISCUSSION

### Connection between Cu nutrition and fatty acid metabolism

During stress adaptation, photosynthetic organisms modify thylakoid membrane fluidity by increasing fatty acid desaturation to maintain photosynthetic performance. In *Chlamydomonas*, Cu limitation triggers remodeling of thylakoid lipids, concomitant with the induction of genes coding for key desaturases such as *SAD*, *FAD5A*, and *FAD6* (Castruita et al., 2011).

Because oleic acid, the primary product of SAD, is a critical component of major lipids in the photosynthetic membrane like MGDG and DGDG, the proper function of SAD is essential for maintaining the integrity of the bioenergetic membrane. Intriguingly, Cu-deficient wt strains exhibit significantly reduced levels of 18:1^Δ9^ in MGDG and DGDG (Figure 3bc). This suggests that the metabolic demand for downstream polyunsaturated products (such as 18:3^Δ9,12,15^) exceeds precursor production, despite the 4-fold, Cu-conditional upregulation of SAD. The increased 18:1^Δ9^ demand is likely the result of metabolic adjustments observed in a Cu deficient cell (Castruita et al., 2011). Those adjustments are likely necessitated by a reduced respiratory rate due to impaired function of the Cu protein COX2b, but also to adjust to subtle changes in the photosynthetic membrane due to the replacement of the Cu protein plastocyanin with the heme containing cytochrome *c*_6_. Furthermore, Cu-deficient cells accumulate triacylglycerols (TAGs) (Figure S2) and (Kropat et al., 2011) and we found that Cu deficient cells still prioritize TAG biosynthesis even when SAD activity is reduced in ami-*sad* mutants (Figure S2).

### FDX5 as an *in vivo* electron donor for SAD

Plant type Ferredoxins (Fds) are small, soluble electron donors to various, Fd dependent client enzymes within the plastid (Fukuyama, 2004). The Chlamydomonas Fd isoform FDX5 has been implicated as electron donor for the Fe hydrogenases based on increased presence in dark anoxic cells (Jacobs et al., 2009; Lambertz et al., 2010), with FDX5 being in stoichiometric excess of the Fe hydrogenase HYDA1 in anaerobic cells (Nikolova et al., 2018). Another study implied FDX5 to be involved in fatty acid metabolism, since FDX5 was shown to interact with fatty acid desaturases FAD6 and Δ4FAD using yeast two-hybrid assays, but changes in the fatty acid profile in *fdx5* mutants were inconsistent with what would have been expected in cells with a loss in either FAD6 or Δ4FAD function (Yang et al., 2015). Direct evidence for physical interaction between FDX5 and the client proteins SAD had not been established prior to this study (Figures 4 and 5b). Although SAD and FDX5 physically interact, the lack of a lipid phenotype in *fdx5* strains, specifically the absence of further 18:1^Δ9^ depletion in our work, contrasts sharply with the dramatic 192-fold induction of *FDX5* transcripts previously reported (Castruita et al., 2011). Despite the magnitude of transcriptional regulation, our immunoblot data suggests that FDX5 is not the predominant Fd isoform in Cu deficient cells, accumulating to just 0.4% of FDX1 levels. This indicates that while FDX5 and SAD physically interact, this interaction does not represent an obligatory bottleneck for thylakoid lipid remodeling in a Cu deficient cell. In Chlamydomonas, the robust constitutive abundance of the major, photosynthetic FDX1 appears sufficient to maintain SAD activity even in the absence of FDX5.

Commonly observed, functional redundancy between Fd isoforms when performing *in vitro* assays suggests that multiple Fd isoforms may share overlapping client networks to ensure metabolic flexibility. The physical association detected by CoIP-MS may reflect a subtle preference, perhaps driven by surface charge complementarities, rather than an absolute requirement. Ultimately, the substantially higher abundance of FDX1, which outnumbers FDX5 by more than two orders of magnitude (Figure 5), likely provides sufficient stoichiometric buffering to compensate for the loss of the specialized FDX5 isoform. To test the limits of this redundancy, future studies should focus on the potential impact that simultaneous repression of both FDX1 and FDX5 has on lipid metabolism, particularly under Cu limitation.

### FDX5: A route to balance carbon metabolism in NADPH rich environments?

FDX5 is predominantly present under two specific growth conditions: Cu deficiency and dark hypoxia (Castruita et al., 2011; Hemschemeier et al., 2013). The physiological commonality between these environments is the generation of high reducing power, where the NADPH/NADP^+^ ratio increases due to metabolic bottlenecks. In Cu deficiency, mitochondrial respiration is impaired by the reduction of the Cu-containing COX2b subunit, for which cells do not possess a Cu-free substitute (Kropat et al., 2015). Similarly, in hypoxic strains, the absence of oxygen shuts down the water-water cycle and other primary electron sinks. In both situations, Chlamydomonas cells induce a genetic program that promotes fermentative pathways like the production of hydrogen, formate, ethanol, or acetate (Mus et al., 2007; Hemschemeier et al., 2013; Yang et al., 2014).

When NADPH levels are saturating, the Ferredoxin-NADP^+^ Reductase (FNR) reaction, which typically transfers electrons from Fd to NADP^+^, can run in reverse. We propose that FDX5 evolved to exploit this high NADPH availability, maintaining reductive power toward specialized metabolic pathways such as lipid desaturation (via SAD) and hydrogen production (via Fe hydrogenase). This hypothesis is supported by our observation that FDX5 interacts directly with both enzymes *in vivo*, consistent with pull down assays reported in work from Subramanian and colleagues (Subramanian et al., 2019).

While FDX5 may be the more efficient electron donor under these specific conditions, FDX1 is appears capable of compensating. Indeed, *in vitro* assays have shown that FDX1 and FDX5 are equally efficient at donating electrons to Fe-hydrogenases (Winkler et al., 2010).

This high degree of functional redundancy between the two Fd isoforms likely explains why *fdx5* mutants exhibit only minor phenotypes at both the physiological and molecular levels. FDX5 does not appear to function as an essential metabolic switch under the conditions tested. Rather, it may function as a fine-tuning mechanism to balance plastid metabolism in cells grown in Cu deficient or anaerobic environments.

## ACKNOWLEDGMENTS

We are thankful to the MSU proteomics core facility and Douglas Whitten for mass-spectrometry based protein identification of immunoprecipitated samples. We thank Sabeeha Merchant for kindly providing Plastocyanin-specific antisera and Chris Jeans and the QB3 Macrolab at UC Berkeley for purification of LbCpf1. This research was primarily funded by the U.S. Department of Energy, Office of Science, Basic Energy Sciences grant towards MSU-PRL (DE-FG02-91ER20021). CB was supported by MSU AgBioResearch. Proteomics work was performed at the Environmental Molecular Sciences Laboratory, a Department of Energy (DOE) Office of Science User Facility sponsored by the Office of Biological and Environmental Research (BER) located at Pacific Northwest National Laboratory (PNNL), proposal ID 60776. Pacific Northwest National Laboratory is operated by Battelle for the DOE under Contract DE-AC05-76RL01830.

## MATERIAL AND METHODS

### Chlamydomonas strains and growth conditions

Chlamydomonas parental strain used for all experiments was CC-425, a cell-wall reduced arginine auxotrophic strain. Cells were grown in tris-acetate-phosphate (TAP) medium prepared with revised trace elements (Kropat et al., 2011) with constant agitation at 120 rpm in a shaker incubator (Multitron, INFORS HT, Annapolis Junction, MD, USA) at 24°C in continuous light (60 µmol m^−2^ s^−1^) (Kropat et al., 2011).

### Generation of Chlamydomonas mutant strains using CRISPR/Cpf1

Chlamydomonas CC-425 was used for transformation with a ribonucleoprotein (RNP) complex consisting of a guide RNA (gRNA) targeting a protospacer adjacent motif (PAM, TTTV) sequence and LbCpf1 as described in Ferenczi et al. (2017); Pham et al. (2022); Strenkert et al. (2024). PAM sequences and gRNAs (Table 1) were selected at the last exon and the last intron of *SAD* and *FDX5* coding sequences to generate HA-tagged strains, respectively (Table S1 and Figure S1 and S5), while the PAM sequence used to generate *fdx5* null mutants were at the second exon of the *FDX5* coding sequence (Table 1 and Figure S6). To generate the gene knock-in of choice, a single-stranded oligodeoxynucleotide (ssODN) containing homology arms to the native locus as well as the respective sequence to be introduced was included in the transformation (Table 1). In brief, a total of 2×10^7^ cells were collected by centrifugation at 2,000*g* for 3 minutes at 21°C. The cell pellet was washed twice with 1 mL of MAX Efficiency Transformation Reagent (Invitrogen). Washed cells were then resuspended in 230 μL of the same solution supplemented with 40 mM sucrose. The cell mixture was heat shocked at 40°C for 30 minutes in a water bath. The RNP complex was assembled by mixing 80 µg of purified LbCpf1 with 2 nmol gRNA (in 10μL) and incubated at 24°C for 30 minutes. Heat-shocked cells were transferred to a 4-mm gap electroporation cuvette. Preassembled RNP complex and HindIII-linearized plasmid containing *ARG7* gene for selection were added along with 4 nmol (10 μL) of the respective ssODN. Electroporation was performed with a Gene Pulser Xcell (Bio-Rad Laboratories, Hercules, CA, USA) with the following setting: 600 V, 50 μF, 200 Ω. After electroporation, the cells were transferred to a 15-mL centrifuge tube containing 4 mL of TAP medium with 40 mM sucrose and incubated overnight in the dark without shaking. On the next day, the cells were centrifuged at 1,160 g, for 3 minutes at 21°C and combined with 60% sterile corn starch resuspended (in TAP with 40 mM sucrose and 0.8% PEG 8000 (w/v)). The cells-starch mixture was then gently spread over TAP agar and incubated under the light for 10-14 days until transformant colonies appear. The colonies were picked and grown on index TAP agar plates for further genotyping.

### Crude DNA extraction and genotyping of transformants

For genotyping of transformants, colony PCR was performed. Chlamydomonas cells were transferred from TAP agar plates and resuspended in 100 μL of TE buffer (10 mM Tris, 1 mM EDTA, pH 8.0). The cell suspension was then heated at 95°C for 10 minutes followed by vigorous vortexing and centrifugation at maximum speed for 5 minutes. The supernatant was used for Polymerase Chain Reaction (PCR) for genotyping using either GoTaq Green Mastermix (Promega, Madison, WI, USA) with the addition of 1 M Betaine or for quantitative PCR (qPCR) with SYBR Green Mastermix (iTAQ mastermix, BioRad) and addition of 1 M Betaine. The primer sequences used for screening the transformants are provided in Supplemental Table 1.

For screening of FDX5-HA strains, PCR products were run on 2% agarose gel with TAE buffer (40 mM Tris-acetate, 1 mM EDTA, pH 8.3). The positive FDX5-HA colonies were selected based on a size-shift of the expected DNA band by 36 bps.

Screening of mutants was done using diagnostic primers that specifically amplify either transgenic or endogenous DNA sequences.

After selection of candidates, DNA sequences were assessed by amplification of the gene edited region with sequencing primers (Supplemental Table 1) and using GoTaq Green Mastermix (Promega, Medison, WI, USA) with the addition of 1 M Betaine. The PCR products were loaded into 1% agarose gel for gel electrophoresis with TAE buffer. The DNA fragment was then extracted using the E.Z.N.A. Gel Extraction Kit (Omega Bio-tek) according to the instructions followed by sanger sequencing.

### Protein isolation

A total of 2×10^7^ cells were collected from cultures at mid-logarithmic phase by centrifugation at 3,000*g* for 3 minutes at 4°C. For equal loading based on protein numbers, cell pellets were resuspended in 100-200 μL of 2x Laemmli’s sample buffer (125 mM Tris-HCl pH 6.8, 20% Glycerol, 4% SDS, 10% β-Mercaptoethanol, and 0.005% Bromophenol blue) and incubated at 65°C for 20 minutes, followed by ice incubation for 2 minutes and kept at −80°C until further use. For experiments that require protein quantification, the cell pellets were resuspended in 50-200 μL of 10 mM sodium phosphate buffer pH 7.0 (4.23 mM NaH_2_PO_4_, and 5.77 mM Na_2_HPO_4_) with or without protease inhibitor cocktail (Roche) and kept at −80°C until further use. Total protein concentration was determined using the Bicinchoninic Acid (BCA) assay (Pierce™ BCA Protein Assay Kits, Thermo Scientific, Waltham, MA, USA). Samples were mixed with 1 volume of 2x Laemmli’s sample buffer and incubated at 65°C for 20 minutes, followed by ice incubation for 2 minutes.

### SDS-PAGE and immuno-detections

Unless noted otherwise, 10 μg total protein was loaded on each acrylamide gel. The protein was separated using SDS-PAGE and blotted onto 0.45 μm nitrocellulose membrane for western blotting using Power Blotter (Invitrogen). The membranes were stained for total proteins with Ponceau S. solution (0.1% Ponceau S (w/v), 5% acetic acid) before incubation with 3% non-fat dried milk (w/v) in PBST (137 mM NaCl, 2.7 mM KCl, 10 mM Na_2_HPO_4,_ 2 mM K_2_HPO_4_, and 0.1% Tween-20) for 1 h. The membranes were then incubated in the primary antibody: anti-HA (1:5.000), anti-SAD (1:1000), anti-Plastocyanin (1:1000, gift from Sabeeha Merchant), anti CF_1_ (1:40.000, gift from Sabeeha Merchant) in 3% non-fat dried milk (w/v) in PBST with constant agitation. The membranes were then washed 4-6 times with PBST and subjected to incubation with a goat anti-rabbit antibody conjugated with alkaline phosphatase at 1:5000 (v/v) in 3% non-fat dried milk (w/v) PBST for 1 h with constant agitation. The membranes were washed as previously described. The alkaline phosphatase signal was detected with its substrate.

### Generation of artificial microRNA based *sad* knock down strains

The artificial microRNA (amiRNA) targeting *SAD* was selected and designed using the WMD3-Web MicroRNA Designer (http://wmd3.weigelworld.org) into pMS692 as described previously (Schmollinger et al., 2010). The target sequence and primers are listed in Supplemental Table 1. The resulting plasmid was linearized with HindIII-HF (New England Biolabs) for transformation into Chlamydomonas CC-425 through glass bead transformation method. In brief, 3×10^7^ cells were collected by centrifugation at 2000 *g* for 3 min The cell pellet was resuspended in 350 μL of TAP with 5% Polyethyleneglycol 8000 (PEG 8000) and transferred to 15 mL centrifuge tube containing 50 μL of sterile glass beads (425-600 μm, Sigma-Aldrich). The cells were vortexed at maximum speed for 15 seconds. The cells were then transferred to a new tube with 4 mL TAP medium with 5% PEG 8000 and incubated overnight in the dark without shaking. On the next day, the cells were centrifuged at 3,000*g* for 3 minutes and resuspended with 650 µL of 60% sterile starch (in TAP with 40 mM sucrose and 1% PEG 8000) before being spread on TAP agar plates. To screen for candidates that had reduced SAD function, cells from agar plates were directly resuspended in 1 mL of 1 M hydrogen chloride (HCl) in methanol. The mixture was overlaid with N_2_ stream, capped and incubated at 80°C for 25 minutes. After incubation, 1 mL of 0.9% NaCl and 150 μL of hexane were added. The mixture was mixed and centrifuged at 3,000*g* for 3 minutes. The hexane (upper) phase was transferred into a GC vial for subsequent analyses of the fatty acid profile.

### RNA extraction and qRT-PCR

A total of 5-8×10^7^ cells were collected by centrifugation at 1610 *g* for 2 minutes at 4°C. The cell pellet was resuspended in 1 mL of TRIzol^®^ Reagent (Invitrogen). Two hundreds μL of chloroform were added, followed by mixing and centrifugation at 13,200 rpm (Eppendorf, 5430 R) for 15 minutes at 4°C. Five hundred μL of the supernatant was transferred to a new tube containing 700 μL of isopropanol. The mixture was incubated on ice for 15 minutes followed by centrifuging at 12,000 rpm (Eppendorf, 5430 R) for 15 minutes at 4°C. The resulting pellet was dried and resuspended in 40 μL of RNase-free water. DNA digestion and RNA clean-up were carried out with the Zymo Clean & Concentrator 5 kit (Zymo Research) by following the instruction manual. cDNA was synthesized with M-MLV Reverse Transcriptase (Invitrogen). qRT-PCR was performed using SYBR Green MasterMix.

Sequences of the FDX5 qRT-PCR sequences are in Supplemental Table 1. The PCR cycle is as followed; 95°C for 10 minutes, followed by 40 cycles of 95°C for 15 seconds and 65°C for 1 minute, 95°C for 15 seconds, 60°C for 1 minute, the temperature is then increased to 95°C at 1%/second, temperature is then reduced to 60°C and hold for 1 minute, and reduced to 4°C for storage.

### Immunoprecipitation followed by Mass spectrometry (IP-MS)

Between 5×10^8^ to 1×10^9^ cells were collected by centrifugation at 1610*g* for 3 min at 4°C. Cell pellets were washed twice with 15 mL of KH buffer (20 mM HEPES-KOH pH 7.2 and 80 mM KCl) and were resuspended in 1 mL of lysis buffer (20 mM HEPES-KOH pH 7.2, 1 mM MgCl_2_, 1 mM KCl, 15 mM NaCl, 1x protease inhibitor cocktail (Roche), and 0.1% triton X-100). The cells were then broken by sonication on ice. Cell debris and intact cells were removed by centrifugation at 1610*g* for 3 min minutes at 4°C. The supernatant was transferred to a new 1.5 mL centrifuge tube and combined with Protein A Sepharose beads coupled with anti- HA antibodies. The protein-bead mixture was incubated with constant rotation at 4°C for 2 hours. The beads were then recovered by centrifugation at 5000 rpm for 20 s at 4°C, washed three times with lysis buffer, washed twice with 10 mM Tris-HCl (pH 7.6). The supernatant was removed, and the beads were frozen for subsequent protein identification by mass spectrometry (MS). For proteolytic digestion, antibody-bound proteins were digested on-bead by washing 3 times using 50mM ammonium bicarbonate. Trypsin, in the same buffer, was then added to the beads at 5ng/µL so that the beads were just submerged in digestion buffer and allowed to incubate at 37°C for 6 hours. The solution was acidified to 1% with trifluoroacetic acid and centrifuged at 14,000 *g*. Peptide supernatant was removed and concentrated by solid phase extraction using StageTips (Rappsilber et al., 2007). Purified peptides eluates were dried by vacuum centrifugation and frozen at −20 C or re-suspended in 2% acetonitrile/0.1%TFA to 20µL. For LC/MS/MS analysis, an injection of 10µL was automatically made using a Thermo (www.thermo.com) EASYnLC 1200 onto a Thermo Acclaim PepMap RSLC 0.1mm x 20mm C18 trapping column and washed for ∼5min with buffer A. Bound peptides were then eluted over 35min onto a Thermo Acclaim PepMap RSLC 0.075mm x 250mm resolving column at a constant flow rate of 300 nL/min with a gradient of 8%B to 22%B from 0min to 19min and 22%B to 40%B from 19min to 24min. After the gradient the column was washed with 90%B for the duration of the run (Buffer A = 99.9% Water/0.1% Formic Acid, Buffer B = 80% Acetonitrile/0.1% Formic Acid/19.9% Water.

Column temperature was maintained at a constant temperature of 50°C using an integrated column oven (PRSO-V2, Sonation GmbH, Biberach, Germany). Eluted peptides were sprayed into a ThermoScientific Q-Exactive HF-X mass spectrometer (www.thermo.com) using a FlexSpray spray ion source. Survey scans were taken in the Orbitrap (60000 resolution, determined at m/z 200) and the top 15 ions in each survey scan are then subjected to automatic higher energy collision induced dissociation (HCD) with fragment spectra acquired at a resolution of 7500. The resulting MS/MS spectra are converted to peak lists using Mascot Distiller, v2.8.5 (www.matrixscience.com) and searched against a protein sequence database containing Chlamydomonas entries based on the most recent *Chlamydomonas reinhardtii* genome sequences (Craig et al., 2023), appended with common laboratory contaminants (downloaded from www.thegpm.org, cRAP project) using the Mascot searching algorithm, v 2.8.3. (Perkins et al., 1999) The Mascot output was then analyzed using Scaffold, v5.3.3 (www.proteomesoftware.com) to probabilistically validate protein identifications. Assignments validated using the Scaffold 1%FDR confidence filter are considered true. Mascot parameters for all databases were as follows: allow up to 2 missed tryptic sites, variable modification of Oxidation of Methionine, peptide tolerance of +/- 10ppm, MS/MS tolerance of 0.02 Da, FDR calculated using randomized database search. Venn diagrams to illustrate interacting proteins that were enriched in IPs from strains expressing HA-fusion proteins, but not in wildtype IPs, were generated using the online tool InteractiVenn (Heberle et al., 2015).

### Lipid analysis

50 mL of Chlamydomonas cells in mid-logarithmic phase cultures were collected by centrifugation at 3,000 *g* for 3 minutes and either directly processed or stored at −80°C. Total lipids were extracted using the modified method from Bligh and Dyer (1959); (Wang and Benning, 2011). The cell pellets were resuspended in 3 mL of methanol, chloroform and formic acid mixture (2:1:0.1 v/v/v). One ml of 0.2 M phosphoric acid (H_3_PO_4_) in 1M potassium chloride (KCl) was added. Samples were mixed and then centrifuged at 3,000*g* for 3 minutes. After phase separation, the lower phase was collected and dried under N_2_ stream. Dried lipid was stored at −20°C. Lipid separation was carried out through thin-layer chromatography (TLC) technique. Dried lipids were dissolved with 150-200 μl of chloroform. Thirty and 5 μl of dissolved lipid was loaded onto a 20×20 cm TLC plate for separation and total lipid control (no separation), respectively. Polar lipids were initially separated with the solvent for polar lipid separation for 5.5 cm from the origin where the samples were loaded. The plate was dried completely before subjected to neutral lipid separation for another 3.5 cm from dye front with the solvent for neutral lipid separation. Solvent systems used in TLC separation are described in Warakanont et al. (2015). TLC-purified lipids were reversibly stained with iodine vapor, scraped and transferred to a glass tube containing 25 μg of 15:0 free fatty acid standard for further conversion to fatty acyl methylesters (FAMEs) as described by Benning and Somerville (1992). FAMEs of TLC-purified lipids and total lipids were dissolved in 70 and 150 μL of hexane, respectively. The FAMEs were then analyzed with a gas chromatography flame ionization detector (GC-FID) on an HP7890 for the ami *sad* samples or on an HP6890 for the *fdx5* samples with a DB-23 column (all from Agilent Technologies, Santa Clara, CA, USA) following this temperature profile; 130°C for 1 min, an increase to 170°C at 6.5°C/min, further increases to 194°C and to 230°C at 8 and 10°C/min, respectively. All samples were run at constant hydrogen (H_2_) flow at 40 mL/min and a split ratio of 1:2.

#### Protein detection by LC-MS/MS

We collected 10^8^ cells by centrifugation at 1,450 *g* at 4°C for 4 min. Cell pellets were processed using a modified S-Trap mini spin column digestion workflow based on the manufacturer’s protocol (ProtiFi, Fairport, NY, USA). Samples were resuspended in 150 μL of preheated 1× lysis buffer consisting of 5% (w/v) SDS and 50 mM triethylammonium bicarbonate (TEAB), supplemented with 5 mM dithiothreitol (DTT). Samples were briefly bath-sonicated for 30 s to bring the material into solution and then incubated at 55 °C for 15 min with shaking at 500 rpm. Lysates were further disrupted by probe sonication for 3 s at 20% amplitude using a 1/16-in. microtip probe attached to a Sonic Dismembrator Model 505 (Fisherbrand). Lysates were clarified by centrifugation at 11,000 × g for 5 min, after which 50 μL of the supernatant was transferred to a fresh tube. The remaining lysate was retained and stored frozen.

Proteins were alkylated by adding 2.5 μL of 400 mM iodoacetamide (IAA; 20 mM final concentration), followed by incubation at room temperature for 15 min in the dark. Samples were then acidified with 5 μL of 12% phosphoric acid and mixed with 350 μL of S-Trap binding buffer consisting of 100 mM TEAB in 90% LC-MS-grade methanol, pH 7.5. The samples were loaded onto S-Trap mini columns and centrifuged until the entire sample volume had passed through the matrix. Columns were washed three times with 400 μL of binding buffer, with the columns rotated between washes, and then transferred to clean collection tubes.

For digestion, 125 μL of digestion buffer containing 15 μg Lys-C (FUJIFILM Wako, Richmond, VA, USA) in 100 mM TEAB was added to each column. Samples were digested at 37 °C for 3 h, after which 20 μg sequencing-grade modified trypsin (Promega, Madison, WI, USA) was added directly to the columns, and digestion was continued overnight at 37 °C.

The following day, peptides were sequentially eluted with 80 μL of 50 mM TEAB in LC-MS-grade water, pH 8.0; 80 μL of 0.2% (v/v) LC-MS-grade formic acid; and 80 μL of 50% (v/v) LC-MS-grade acetonitrile in 0.2% (v/v) LC-MS-grade formic acid. Columns were centrifuged at 10,000 × g for 1 min after each elution step. Eluted peptides were dried in a SpeedVac, desalted using Waters Oasis tC18 100 mg SPE cartridges (Waters, Milford, MA, USA), and resuspended for LC-MS analysis. Samples were loaded onto LC columns with 0.05% formic acid in water and eluted in 0.05% formic acid in ACN over 100 min. Twelve high resolution (17.5K nominal resolution) data-dependent MS/MS scans were recorded in centroid mode for each survey MS scan (35K nominal resolution) using normalized collision energy of 30, isolation width of 2.0 m/z, and rolling exclusion window lasting 30 seconds before previously fragmented signals are eligible for re-analysis. Unassigned charge and singly charged precursor ions were ignored.

#### MS/MS data search and analysis

The MS/MS spectra from all LC-MS/MS datasets were converted to ASCII text (.dta format) using MSConvert (http://proteowizard.sourceforge.net/tools/msconvert.html) which precisely assigns the charge and parent mass values to an MS/MS spectrum as well as converting them to centroid. The data files were then interrogated via target-decoy approach (Elias and Gygi, 2010) using MSGFPlus (Kim and Pevzner, 2014) using a +/- 20 ppm parent mass tolerance, partial tryptic enzyme settings, and a variable posttranslational modification of oxidized Methionine. All MS/MS search results for each dataset were collated into tab separated ASCII text files listing the best scoring identification for each spectrum. MS/MS search results for each biological replicate were collated to a tab delimited text file using in-house program MAGE Extractor. These results were then imported into a SQL Server (Microsoft, Redmond, WA) database and filtered to 1% FDR by adjusting the Q-Value provided by MSGF+. The in-house program MASIC (https://omics.pnl.gov/software/masic, 53) was run on each dataset, which provides ion statistics and general information extracted from the instrument binary rawfile. Specifically, both the maximum observed MS ion count for each MS/MS spectrum’s precursor signal (termed PeakMaxIntensity) and a selected ion chromatogram (SIC, termed StatMomentsArea) are provided for subsequent peptide quantitation. The results from all datasets’ MASIC output were collated using the in-house program MAGE File Processor, with a tab delimited text file listing all relevant data as the result. These results were also imported into SQL Server and connected to the filtered MSGF+ results via Dataset_ID (internal to PNNL’s data management system), and relevant MS/MS scan number. Unique peptide sequences (with post-translational modifications counted as separate sequences) were grouped per dataset with observation counts and maximum PeakMaxIntensity values provided. This data was then pivoted to provide a crosstab of peptides, with protein information carried through the query. The crosstab was then imported into InfernoRDN (an implementation of the R statistical package, https://omics.pnl.gov/software/infernordn), Log_2_ transformed and mean central tendency normalized (boxplot alignment). The normalized peptide abundance values were then exported into Excel (Microsoft), anti-logged, and proteins grouped with values summed using the Pivot Table function. Most downstream analyses were carried out in R, including the determination of Euclidean distances (stats package) and hierarchical clustering (stats package, using the ward.D2 algorithm). Graphing was performed in R (ggplot2, pheatmap packages) and further modified in Adobe Illustrator. The mass spectrometry proteomics raw data will be deposited after publication.

