## Supplemental Figures and Legends for "SAD-dependent thylakoid lipid desaturation and FDX5-associated electron transfer during copper deficiency in *Chlamydomonas reinhardtii*"

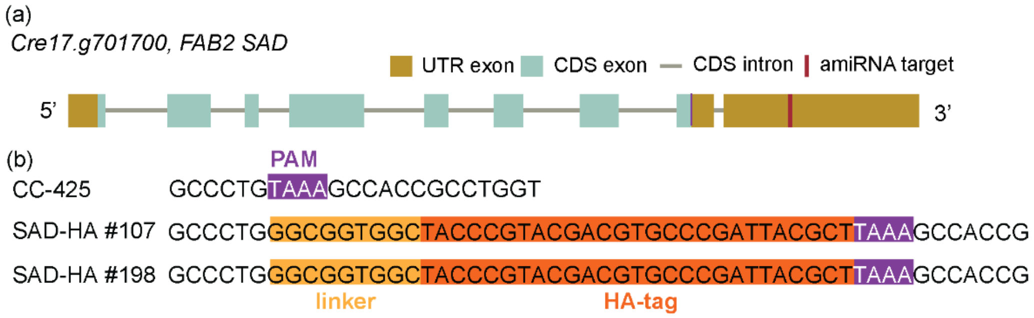


**Supplemental Figure 1. *SAD*/*FAB2* gene model and amiRNA target site/CRISPR knock-in approach.** Diagram of the *FAB2* locus showing the PAM (magenta) and amiRNA target site (red) used to generate SAD-HA knock-in strains using Cpf1 or sad-ami mutants, respectively. A short linker (yellow) and the sequence encoding the HA epitope (dark orange) were inserted in-frame before the stop codon of *FAB2*, yielding strains SAD-HA #107 and SAD-HA #198.

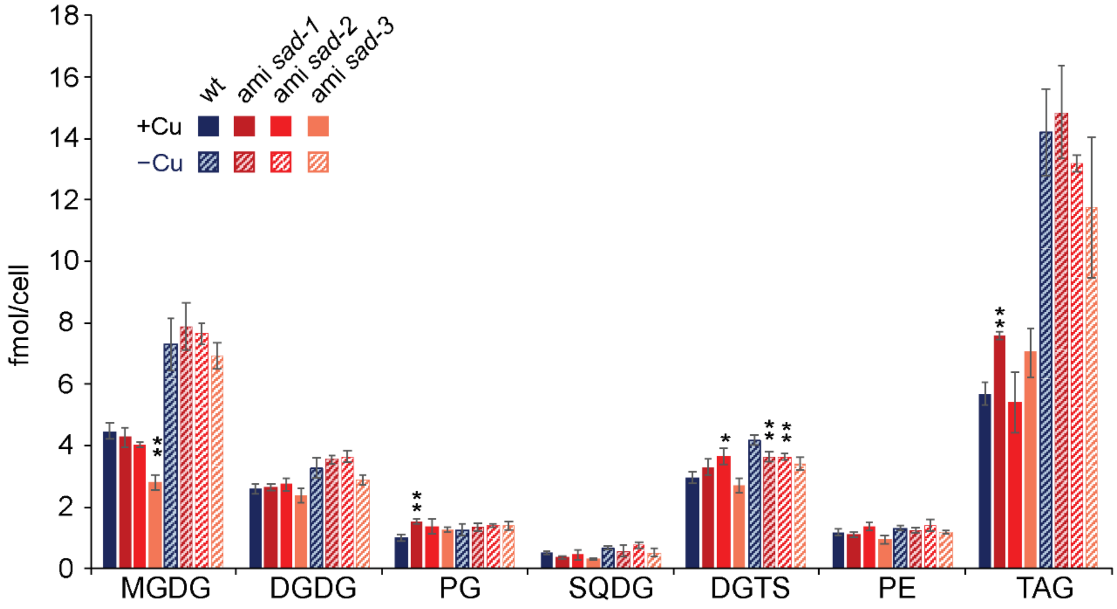


**Supplemental Figure 2. Lipid abundance of ami *sad* strains under Cu replete (+Cu) and deplete (-Cu) conditions.** Abbreviations: DGDG; digalactosyldiacylglycerol, DGTS; diacylglyceryl-N,N,N-trimethylhomoserine, MGDG; monogalactosyldiacylglycerol, PE; phosphatidylethanolamine, PG; phosphatidylglycerol, and TAG; triacylglycerol.

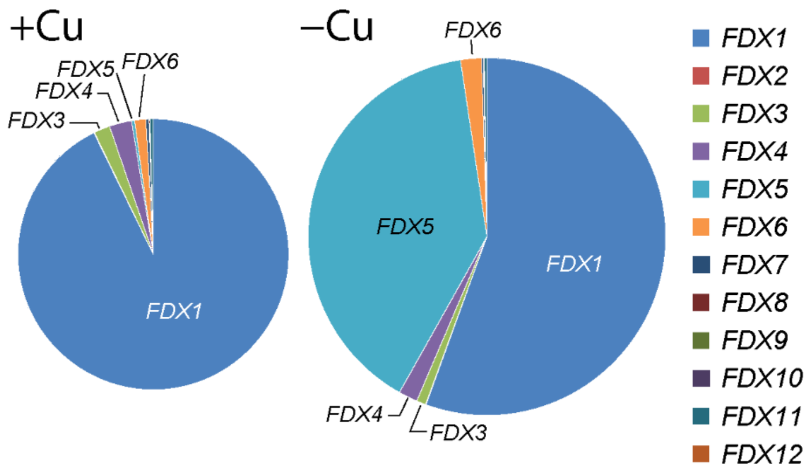


**Supplemental Figure 3. Relative expression of ferredoxin isoforms (*FDX1-12*) in response to changes in the Cu nutritional status.** Shown are *FDX* transcript abundances in cells that were grown in TAP medium with (+Cu) or without Cu supplementation (-Cu) as indicated. The diagrams were drawn to scale of the FPKM abundance from RNA-Seq data from Castruita et al. (2011).

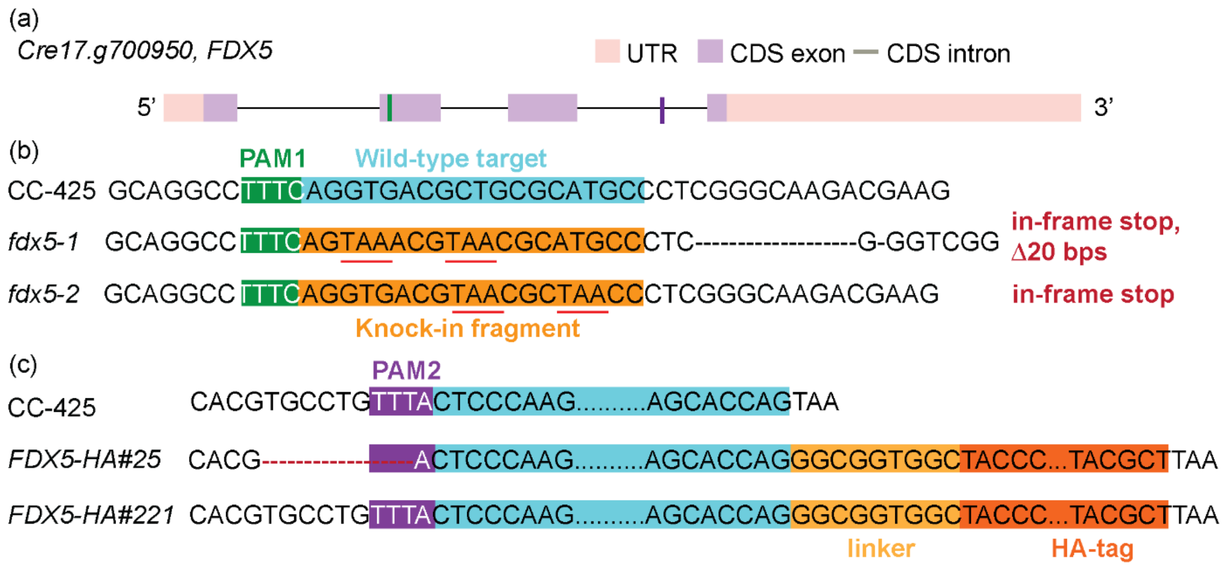


**Supplemental Figure 4. *FDX5* Gene model and CRISPR approach.** Diagram of the *FDX5* locus showing the palindrome-adjacent motifs (PAMs) and the target sites used to generate knockout mutant strains (green) or HA knock-in strains (magenta) via CRISPR-mediated gene editing. Stop codons were introduced in-frame in the *fdx5-1* and *fdx5-2* mutants and are highlighted in red font; a short linker (yellow) was added upstream of the sequence encoding the HA epitope (dark orange) introduced before the native *FDX5* stop codon.

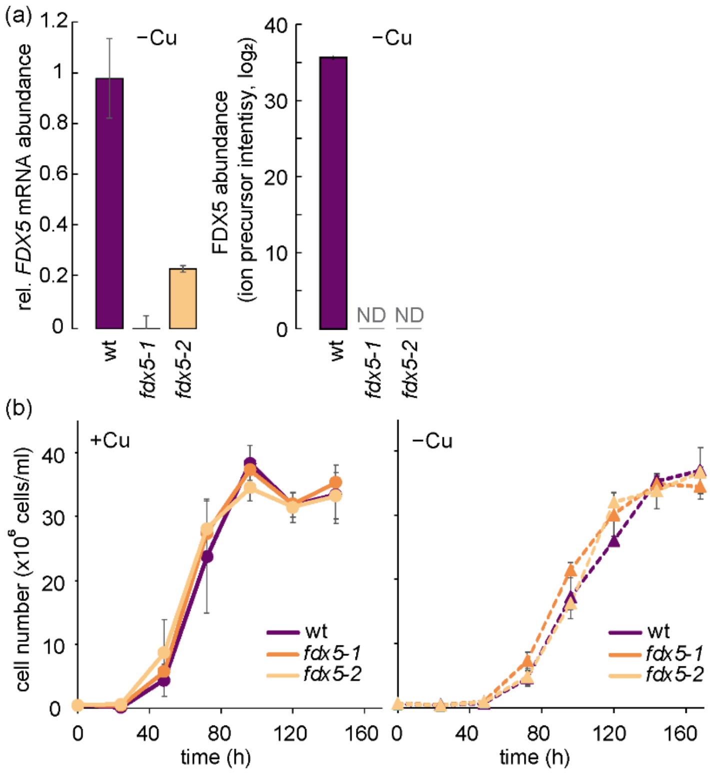


**Supplemental Figure 5. Mutants in *fdx5* are phenotypically asymptomatic independent of the Cu nutritional status.** (a) *FDX5* transcript level and FDX5 protein abundance in *fdx5-1* and *fdx5-2* mutants compared to that of wt based on RT-qPCR analysis and proteomics, respectively. Cells were grown in medium without Cu supplementation. (b) Growth curves of *fdx5-1* and *fdx5-2* mutants and wt under copper replete (+Cu) and deficient (-Cu) conditions. Cells were counted every 24 hours using a CoulterCounter. Shown are averages and STDEV of three independent cultures.

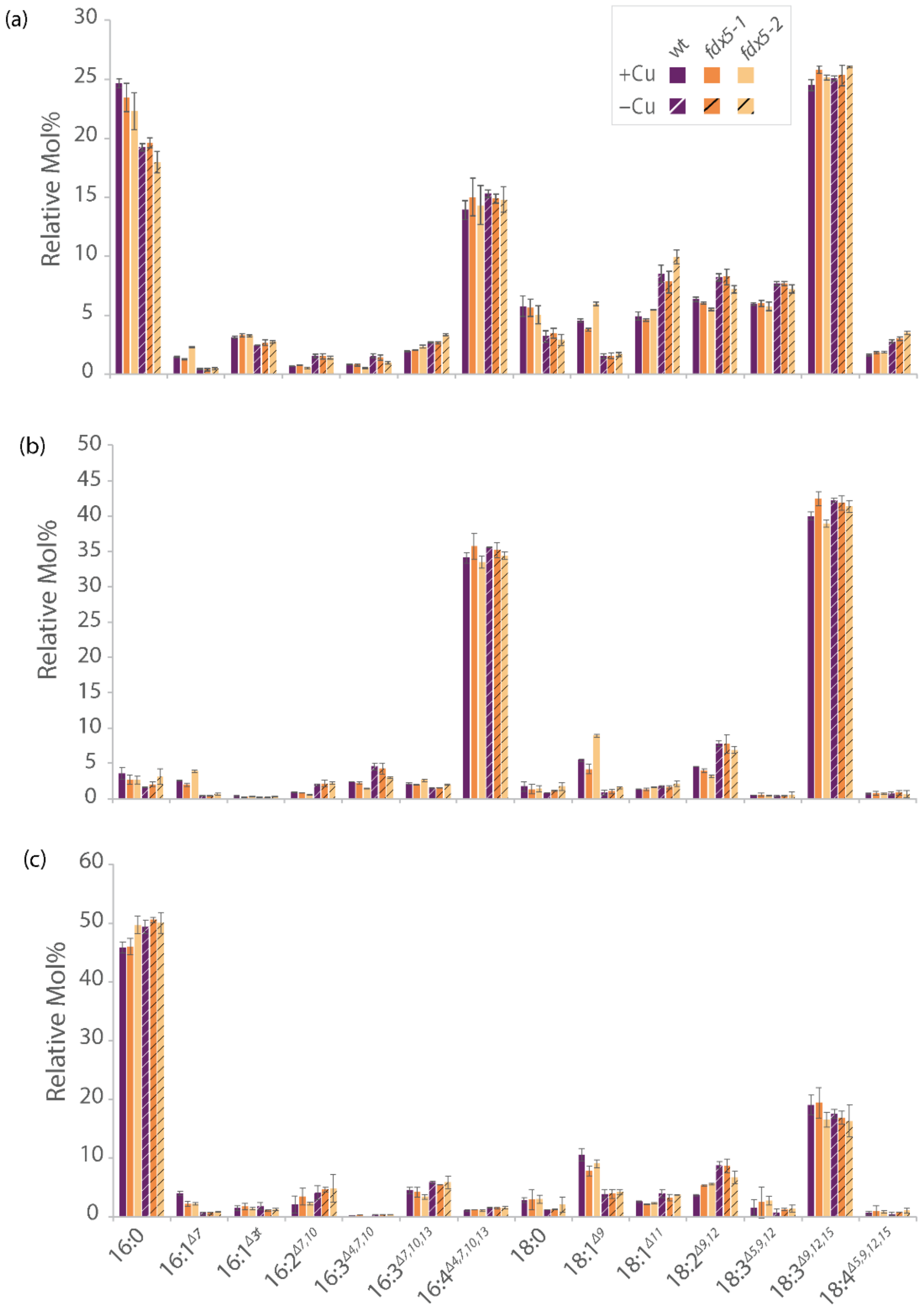


**Supplemental Figure 6. Lipid profile is unchanged in *fdx5* mutants.** Acyl group profile of total lipids (a), monogalactosyldiacylglycerol (MGDG, b) and digalactosyldiacylglycerol (DGDG,c) of the wt and *fdx5* mutants during mid-logarithmic phase grown in different copper (+Cu and -Cu) conditions. The bars show fatty acid composition in mol % and indicate the means (±SD) of three independent experiments. Standard nomenclature for fatty acids is used, and is indicated below the *x*-axis in (c): number of carbons:number of double bonds, with position of double bonds indicated counting from the carboxyl end. A single asterisk (*) and a stacked double asterisk (one * above another *) indicate statistically significant differences from the wild type with $p \leq 0.05$ and $p \leq 0.01$, respectively, based on two-tailed Student's *t*-tests assuming equal variances. Statistical analysis was performed as described in Figure 3.

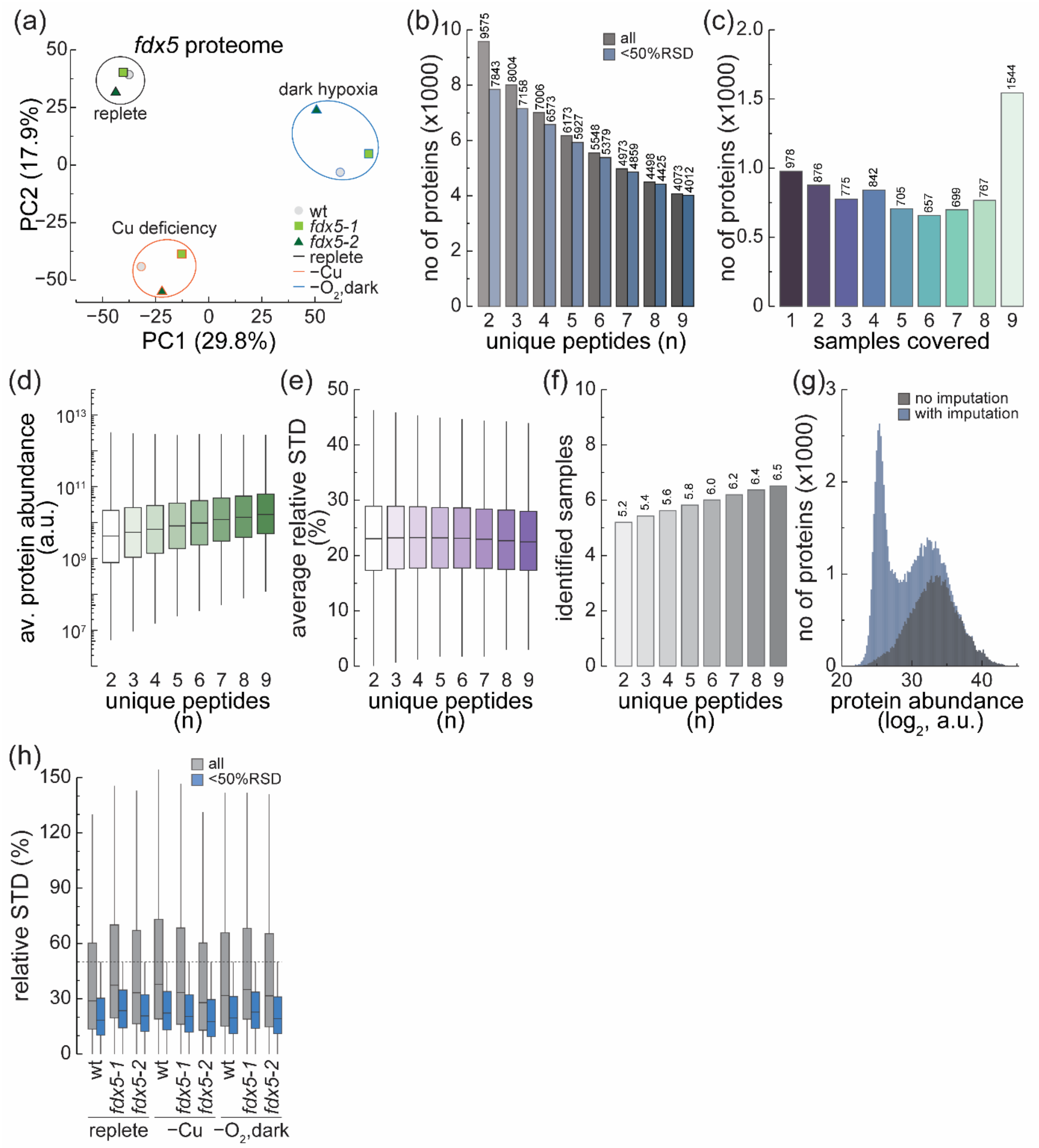


**Supplemental Figure 7. Proteomic changes in *fdx5* mutants grown under Cu deficiency or dark hypoxia.** (a) Principal component analysis (PCA) of the Chlamydomonas proteome of wt (grey circle), *fdx5-1* (green square) and *fdx5-2* mutants (blue triangle), collected under replete, Cu deficiency or dark hypoxia. (b) Proteins identified (No.) dependent on the number of unique peptides required for a protein to be considered identified (grey fill). Blue bars identify only proteins also requiring a relative standard deviation below 50% between replicates. (c) Dataset coverage, number of proteins identified per number of samples the protein was identified in (No.). (d, e, f) Relationship between the minimum unique-peptide requirement and (d) average abundance across all samples (a.u.), (e) average relative SD (RSD) between replicates (%), and (f) average number of samples the protein was identified in. (g) $log_2$-transformed protein abundance distribution of all data without imputation (grey bars) and after imputation of missing data points (blue bars). (h) RSD (%) distribution in samples. Shown are all identified proteins (grey fill), or after filtering for consistency between replicates.

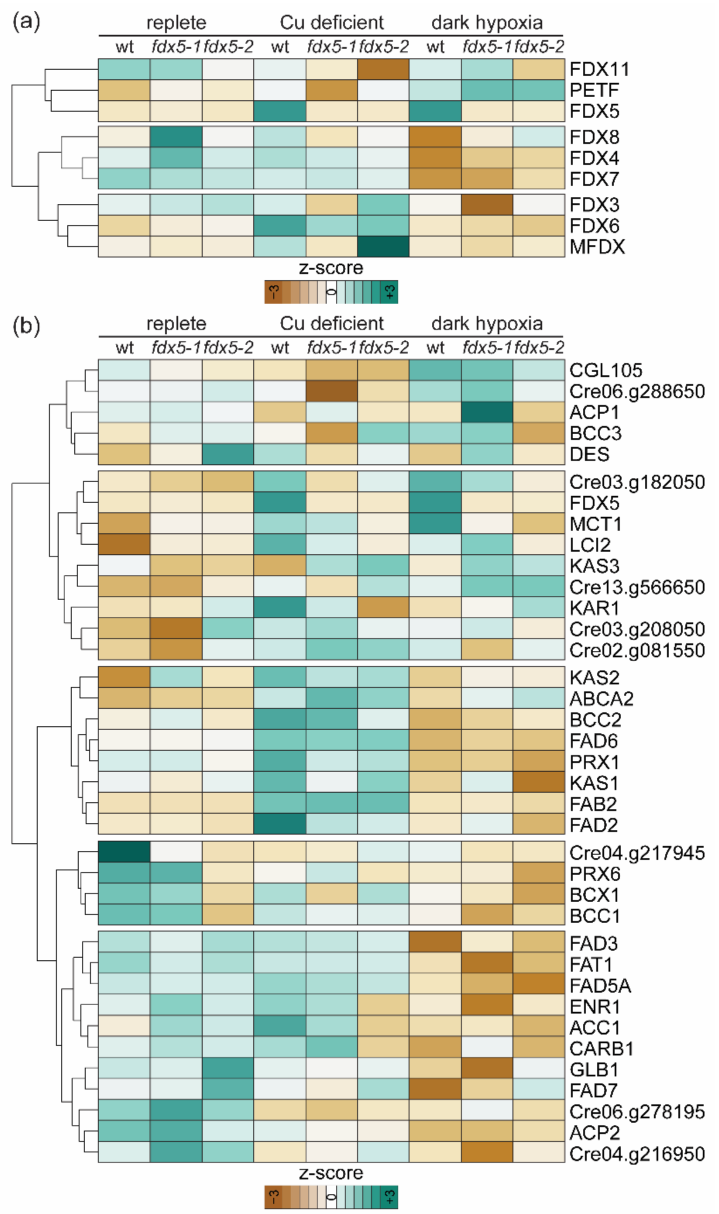


**Supplemental Figure 8. Abundance changes of ferredoxins and proteins involved in lipid synthesis in *fdx5* mutants grown under Cu deficiency or dark hypoxia.** (a) 9 ferredoxins were identified in the proteomics dataset, shown is a heatmap of protein abundances (z-score) over all samples. Ferredoxins were hierarchically clustered based on their response to the stimuli. (b) Expression changes (z-score) of proteins involved in fatty acid biosynthesis.
